# Evolution of Enhancers through Duplication

**DOI:** 10.1101/2025.04.09.648062

**Authors:** Devika Singh, Taylor Hoyt, Soojin V. Yi

## Abstract

Duplication is appreciated as a key source of genomic novelty. In this study, we examined putative enhancers in the human genome to investigate how duplication influences the evolution of these enhancers. We strictly identified a set of 3,336 confident duplicate enhancer pairs and examined their genomic and evolutionary features. We found that, compared to non-duplicated enhancers, duplicated enhancers tend to be longer, more pleiotropic, closer to genes, and harbor greater numbers and more diverse groups of transcription factor binding motifs. These attributes were more pronounced for evolutionarily older duplicate enhancers. Therefore, the regulatory potentials of enhancers may facilitate evolutionary maintenance of duplicated enhancers. Utilizing chimpanzee and rhesus macaque as outgroups, we found that between 30-40% of the examined duplicate enhancers exhibit evidence of asymmetric sequence evolution. Notably, the majority of the “accelerating enhancers” in such pairs gained enhancer activities in novel tissues, particularly in immune-related tissues. Moreover, the accelerating enhancers tended to harbor transcription factor binding motifs previously implicated in human evolution, and enriched in associations with immune functions and stress responses. These findings indicate that duplication may contribute to the proliferation of highly pleiotropic enhancers, as well as gaining novel enhancer activities, and contribute to rapid evolution of the immune system.

## Introduction

Genomic duplication events, encompassing small-scale sequence duplications to whole genome duplication, are canonical sources for the raw materials used for the evolution of functional elements of the genome (Ohno 1970). For example, the frequency and consequences of gene duplications have been extensively studied. Previous works demonstrate that, although gene duplications occur at a high frequency, the vast majority of these redundant regions lose functionality rapidly due to the accumulation of degenerative mutations in a process called non-functionalization ((Innan and Kondrashov 2010; Lynch and Conery 2000) and references therein). On the other hand, duplicate genes may be retained via alternative evolutionary trajectories including neofunctionalization or subfunctionalization. Neofunctionalization refers to the cases when one gene copy retains the ancestral function while the other gains a novel function through the acquisition of an advantageous mutation and subsequent positive selection (Force, et al. 1999; Ohno 1970). In contrast, subfunctionalization refers to instances when degenerative mutations accumulate in both copies of the duplicated gene, but the ancestral gene function is maintained by the combined dosage of the duplicate pair (Lynch and Force 2000). Both of these scenarios demonstrate critical pathways for the expansion of novel gene functions which can increase the functional diversity of the genome. In fact, it is estimated that 15-50% of all human genes have originated via duplication events (Acharya and Ghosh 2016; Keller and Yi 2014; Li, et al. 2001; Park and Makova 2009).

Functional evolution through sequence duplication could also contribute to diversification of non-coding *cis*-regulatory elements (reviewed in (Long, et al. 2016)). Chief among these elements are enhancers, short DNA sequences that control the precise context- and time-dependent expression of genes (Banerji, et al. 1981; Lettice, et al. 2014; Long, et al. 2016). Specific instances of the retention of functional duplicated enhancers have been reported such as the two hepatic control regions driving the expression of the human apolipoprotein (apo) E genes (Allan, et al. 1995; Goode, et al. 2011). Duplicated enhancers are also associated with diverse abnormal phenotypes in humans such as Keratolytic winter erythema (KWE), bilateral concha-type microtia, disorders of sex development (Croft, et al. 2018; Ngcungcu, et al. 2017; Si, et al. 2020). However, a comprehensive, global analysis of how duplicate enhancers evolve has yet to be conducted.

Several factors may influence the differences in evolutionary forces acting on duplications in genic regions compared and those in enhancer regions. Enhancers are organized in a many-to-one interaction structure where there are many more enhancers than genes (ENCODE_consortium 2012; Singh and Yi 2021). This redundancy helps maintain the robust and stable expression of target genes by acting as a buffer, given frequent fluctuations in transcription factor inputs and deleterious mutations in any regulatory regions (Huh, et al. 2018; Kvon, et al. 2021; Osterwalder, et al. 2018; Waymack, et al. 2020). Many enhancers are also highly tissue-specific (Singh and Yi 2021), which collectively suggests that this reduction in effect size compared to a gene may alleviate the selection pressure or evolutionary constraint on an individual enhancer (Sabarís, et al. 2019). On average, enhancers are also shorter than genes and can readily evolve function from ancestral regulatory sequences, which reduces the evolutionary barrier for *de novo* enhancer formation either through spontaneous emergence or exaptation (Fong and Capra 2021; Rebeiz, et al. 2011; Villar, et al. 2015). Therefore, enhancers may have alternative trajectories to evolve novel functions and are subject to less selective constraints associated with the redundant copies.

Here, we aim to examine how duplication and subsequent functional diversification may have occurred over the course of evolution of enhancers in the human genome. We have previously (Singh and Yi 2021) curated and characterized a large dataset of putative enhancers from the Roadmap Epigenomics Mapping Consortium (Roadmap Epigenomics, et al. 2015) across 23 diverse human tissues. A notable result from this study was the identification of a rare (<1%) subset of highly pleiotropic enhancers with an increased effect size in terms of breadth of activity and number of regulated target genes. In this study, we utilize these annotations to determine the frequency of duplicate enhancer maintenance as well as enhancer features which may determine their retention over evolutionary time in the human genome.

## Results

### Distinctive Genomic Features of Duplicate Enhancers

For our analyses, we utilized the curated set of enhancers generated in Singh and Yi (Singh and Yi 2021) from NIH’s Roadmap Epigenomics Mapping Consortium (Roadmap Epigenomics Consortium, et al. 2015). Beginning with 646,419 unique putative enhancers identified from chromatin-state characterization (Abascal, et al. 2020) across 23 human tissues, we performed an all-by-all BLAST coupled with stringent filtration criteria for repetitive content and overlap coverage (see Methods) to identify candidate duplicated enhancer families. We further performed single-linkage clustering (SLC) within duplication groups to identify duplicate enhancer pairs based on evolutionary divergence measured by Kimura’s two-parameter (K2P) model (Kimura 1980). In total, we generated a dataset of 3,336 candidate duplicate pairs with a mean K2P distance of 0.21 (**Supplementary Figure 1**). We consider these as a conservative and strictly defined set of duplicate enhancers that we utilized for downstream analyses.

Given that these duplicate enhancers encompass a small subset of the total number of putative enhancers, we were interested in exploring the genomic characteristics contributing to their continued maintenance over evolutionary time. We specifically considered the relative enrichment of six attributes of duplicated enhancers compared to control groups of length-matched non-duplicated enhancers acting as the genomic background. As we previously defined (Singh and Yi 2021), the degree of “enhancer pleiotropy,” refers to the number of tissues in which a region exhibits an enhancer chromatin state and is consequently considered active. As such, low enhancer pleiotropy values indicate that the corresponding enhancers act in a tissue-specific manner while high values imply that the enhancers are broadly active in multiple tissues. We found that duplicate enhancers are significantly longer and more pleiotropic than non-duplicate control enhancers (**Supplementary Table 1, Figure 1a, b**). In addition, duplicate enhancers are in closer proximity to genes and linked to a greater number of target genes than control enhancers (**Supplementary Table 1, Figure 1c, d**). Finally, we found that duplicate enhancers harbor a significantly greater number and more diverse groups of transcription factor (TF) binding motifs compared to control regions (**Supplementary Table 1, Figure 1e, f**).

**Figure 1.**
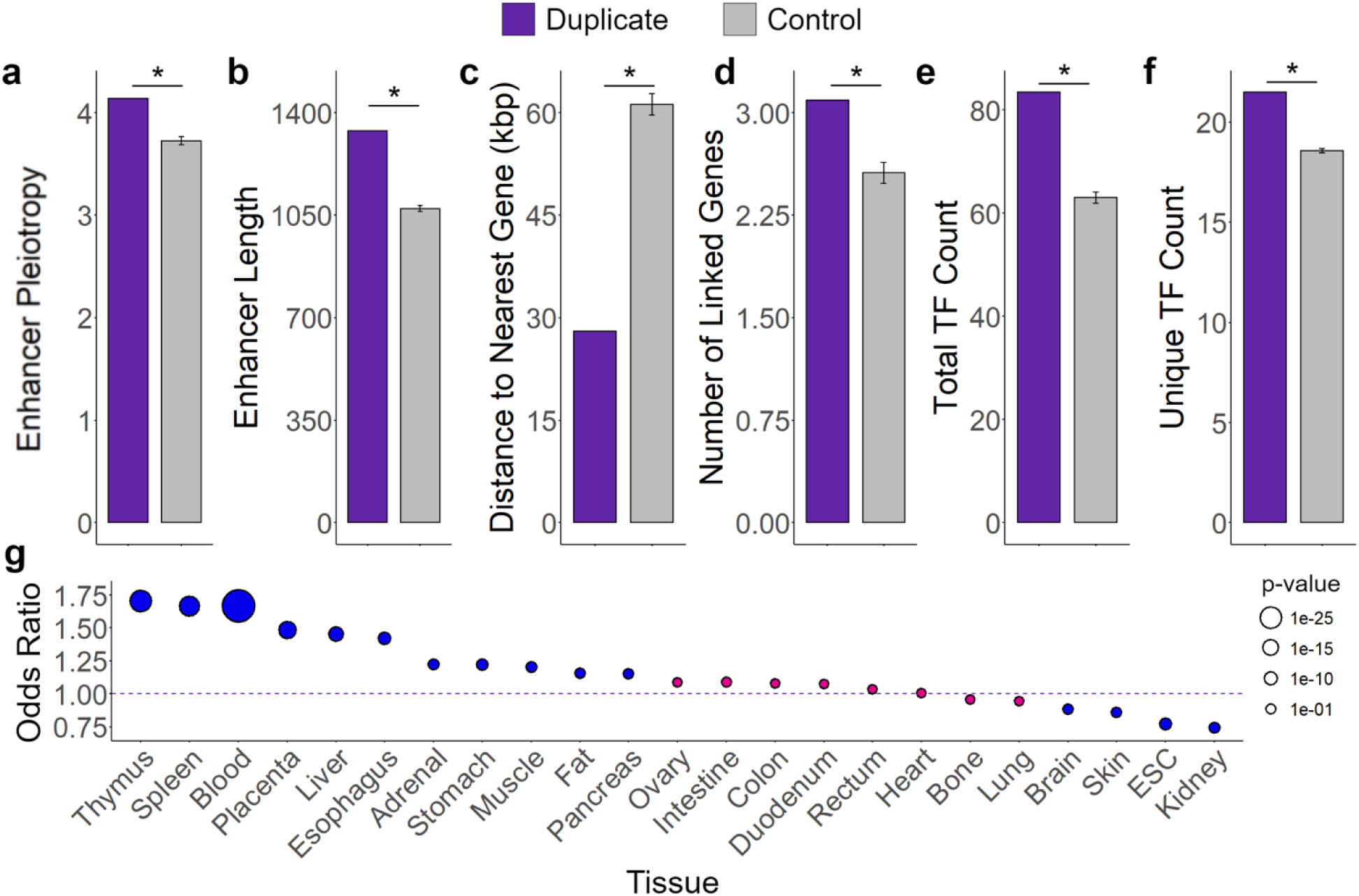
Genomic characteristics of duplicate enhancers. Enrichment of six genomic attributes of duplicate enhancers, (**a**) enhancer pleiotropy, (**b**) enhancer length (bp), (**c**) distance to nearest gene (kbp), (**d**) number of linked target genes per enhancer (**e**) total number of transcription factors (TFs) per enhancer, and (**f**) total number of unique TFs (representing TF diversity) per enhancer, compared to those of length-matched non-duplicate control enhancers. For all attributes (a-f), p < 0.001 (illustrated as *) based on 1,000 bootstraps. Error bars indicate standard deviation. (**g**) Enrichment of duplicate enhancers in all surveyed tissues compared to length-matched non-duplicate control enhancers. Odds ratio and p-value are reported from Fisher’s Exact Test considering the occurrence of duplicate enhancers active or not active in each tissue compared to the expected pattern from the control enhancers.

We further investigated which, if any, of the 23 tissues analyzed were significantly enriched or depleted of duplicated enhancers compared to the control enhancers. We show that 11 out of 23 tissues (47.8%) are significantly enriched for duplicated enhancers in both datasets (**Figure 1g, Supplementary Table 2**). This subset notably includes members of the primary and secondary lymphoid organs, the thymus and spleen as well as blood cells. The brain, skin, embryonic stem cells, and kidney samples were consistently, and significantly, depleted of duplicated enhancers compared to the control datasets (odds ratio < 1, p ≤ 0.002 Fisher’s Exact Test). These observations indicate that duplicate enhancers harbor specific genomic traits and tend to be used in subsets of immune-related tissues.

### The relative age of duplicated enhancers is predictive of regulatory potential

Given that duplicate enhancers show a wide range of sequence divergence between pairs (**Supplementary Figure 1**), we hypothesized that characteristics affecting the regulatory potential of duplicated enhancers could be a factor in their preservation over evolutionary time. To test this hypothesis, we assigned relative ages to each enhancer pair based on their pairwise K2P distances, where smaller distances between pairs imply a more recent duplication while larger distances are indicative of an older duplication. We then binned the duplicate pairs evenly between the minimum and maximum evolutionary distance such that no bin contained less than 10 data points and examined the variation of the aforementioned enhancer attributes (**Figure 2**).

**Figure 2.**
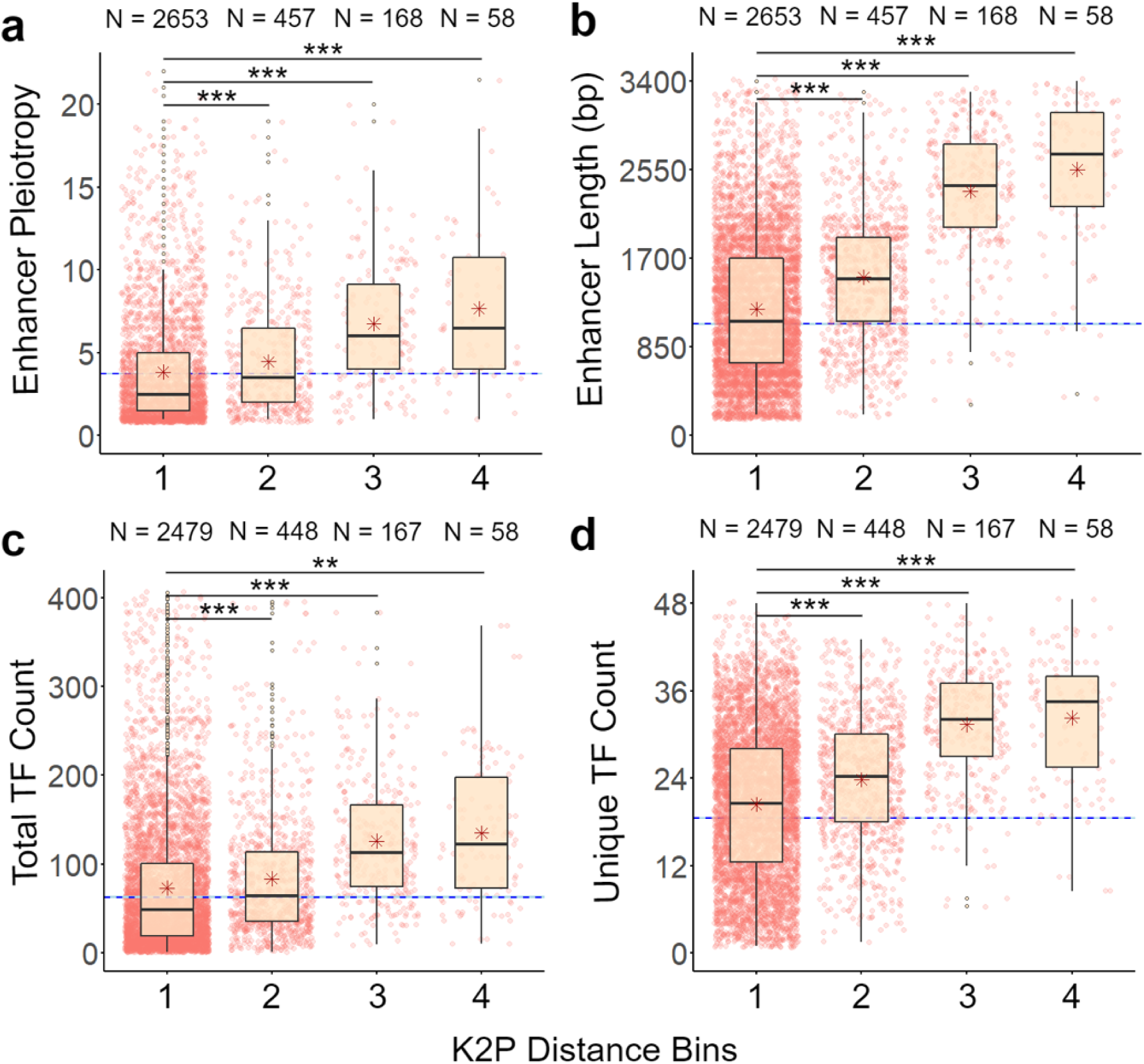
Correlation between relative age of duplicate enhancers and genomic characteristics. Distribution of (**a**) enhancer pleiotropy, (**b**) enhancer length (bp), (**c**) total number of transcription factors (TFs) per enhancer, and (**d**) total number of unique TFs (representing TF diversity) per enhancer for all duplicate enhancers divided into bins by their K2P distance. The bins were evenly distanced between the minimum and maximum K2P of the total duplicate enhancer set. The K2P ranges within each bin are as follows: Bin 1 K2P = 0-0.33, Bin 2 K2P = 0.33-0.67, Bin 3 K2P = 0.67-1.00, and Bin 4 K2p > 1. The total numbers of duplicate enhancers per bin are reported along with p-values from Mann-Whitney U test (*** indicates p < 2.2 × 10-16 and ** indicates p < 0.001). The mean values of length-matched non-duplicate control enhancers are shown as a horizontal dashed line on each plot.

Enhancer pleiotropy, length, and transcription factor count and diversity all increase with relative ages of duplicate pairs such that the “youngest” (K2P distance ≤ 0.33) duplicates are consistently and significantly less pleiotropic, shorter, and harbor fewer transcription factors than the “oldest” (K2P distance > 1.00) duplicate enhancers (p ≤ 0.05 for all attributes, Mann–Whitney *U* test). Indeed, compared to the mean attribute values of the control distributions, the “oldest” duplicates show a larger and more significant deviation than the “youngest” duplicates (**Figure 2** and **Supplementary Table 3**). These results suggest that more pleiotropic, longer enhancers harboring large numbers of transcription factors may be preferentially maintained during evolution.

### Signatures of asymmetric evolution in recently duplicated enhancers

The above analyses indicate that regulatory attributes of enhancer sequences are associated with instances of the evolutionary retention of duplicate enhancers. If the gain and loss of regulatory attributes are related to changes at the sequence level, we may be able to detect the associated signal at the level of sequence evolution. In other words, we can determine whether duplicate enhancers undergo accelerated sequence evolution, which would be expected if the gain of a specific regulatory attribute is driven by positive selection. In the following section we utilized two non-human primate (NHP) genomes (rhesus macaque and chimpanzee genomes) to further examine the fates of duplicate enhancers in relation to their sequence evolution.

To quantify duplicate enhancer sequence evolution, we first employed a sequence homology search to identify orthologous regions in both non-human primate (NHP) genomes (summarized in **Figure 3a**). Briefly, we used a reciprocal best BLAST hit (RBBH) approach to identify single-copy orthologous regions independently in the rhesus macaque and chimpanzee genomes. For the subset of duplicate enhancers with single-copy orthologous regions in both NHP outgroups, we surmised these enhancers likely resulted from a duplication event following the divergence of the human-chimpanzee lineages. Due to the difficulty of aligning non-coding genomic regions of a more distantly related outgroup, duplicate enhancers with single-copy orthologous regions in the macaque lineage may include instances of a loss of duplication in the macaque genome rather than a gain of duplication in the human genome. However, these regions are still informative in analyzing signatures of sequence evolution within human duplicate pairs. Furthermore, we found that only 2.1% (69/329) of the duplicate enhancers with single-copy orthologous regions in the chimpanzee exhibited ‘loss’ in the chimpanzee genome (**Supplementary Table 4**). It stands to reason that most duplicate pairs with single copy orthologous regions in the rhesus macaque genome likely result from duplication events following the human-rhesus macaque divergence. In total, we report 738 and 260 duplicate enhancer pairs with rhesus macaque and chimpanzee as outgroups, respectively (**Table 1**).

**Figure 3.**
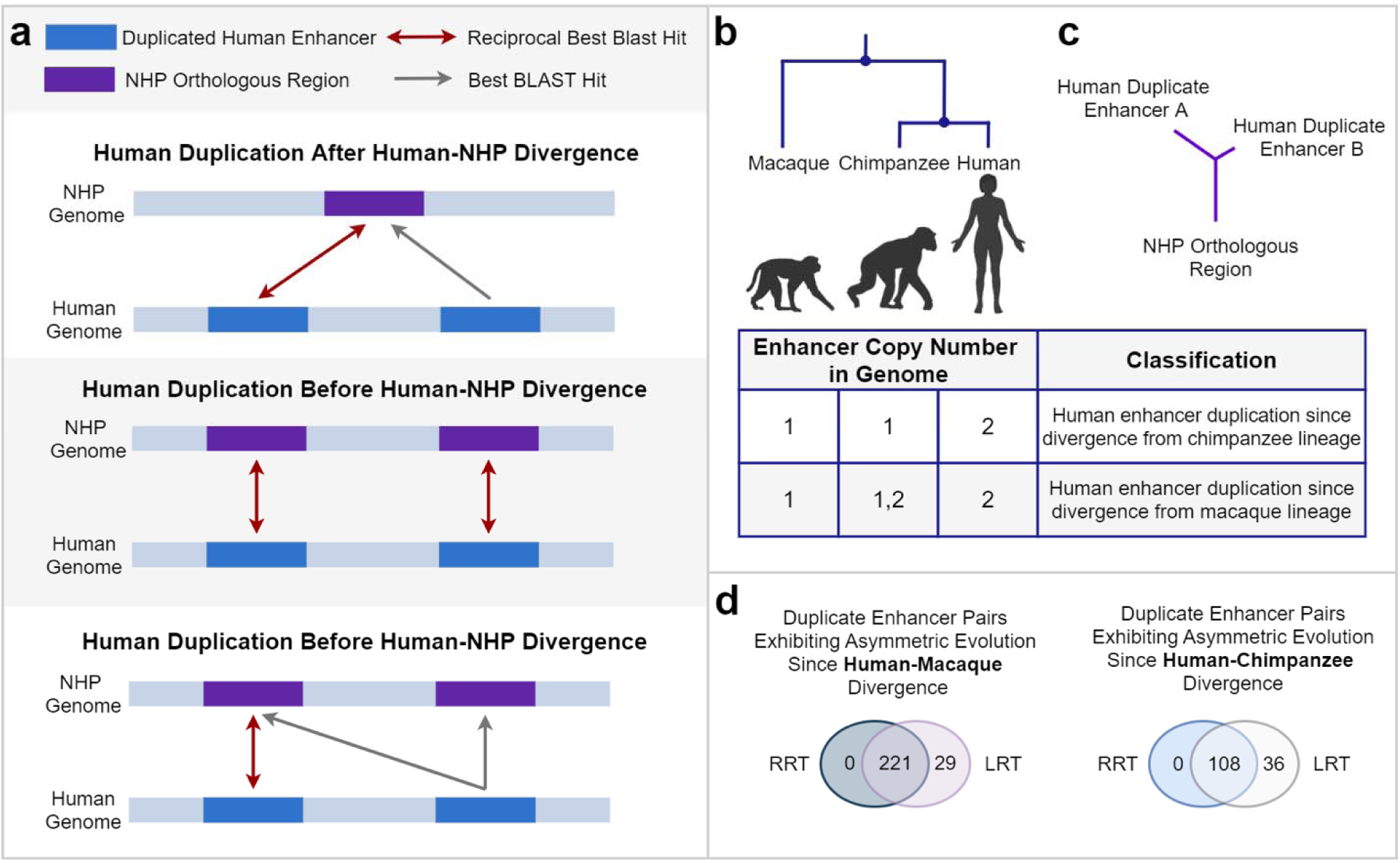
Asymmetric evolution of duplicate enhancers. (**a**) Schematic representation of the reciprocal best BLAST hit (RBBH) strategy to identify orthologous regions in non-human primate (NHP) genomes. Duplicate enhancers with single-copy orthologous regions in one or both NHP genome were identified if one enhancer within a pair was reciprocally the “best hit” for a NHP region while the other enhancer uniquely mapped to the same region. (**b**) Classification scheme for duplicate enhancers with single-copy orthologous regions in one or both NPH genome. (**c**) Representative phylogenic tree of a duplicate enhancer pair in which one enhancer (enhancer A) exhibits accelerated evolution relative to its mate (enhancer B). For these analyses, the NHP orthologous region is used as the outgroup sequence. (**d**) Total number of duplicate enhancers identified as exhibiting significant accelerated evolution in a Relative Rate Test (RRT) and a Likelihood Ratio Test (LRT) utilizing either the chimpanzee or rhesus macaque as the outgroup.

**Table 1.** Duplicate enhancer pairs exhibiting signatures of asymmetric evolution. Duplicate pairs (DPs) with single-copy orthologous sequences in either rhesus macaque or chimpanzee genomes are reported and utilized as outgroup sequences to identify significant asymmetric enhancer sequence evolution.

| Total | Total DP | Asymmetric DP | Total DP | Asymmetric DP |
| --- | --- | --- | --- | --- |
| Duplicate | Rhesus Macaque | Rhesus Macaque | Chimpanzee | Chimpanzee |
| Pairs (DP) | Outgroup | Outgroup | Outgroup | Outgroup |
| 3,336 | 738 | 221 (29.9 %) | 260 | 108 (41.5 %) |

To test for instances of accelerated sequence evolution of enhancers within recently duplicated pairs (**Figure 3c**), we utilized the baseml module from PAML (Yang 2007) which compares the maximum likelihoods of different evolutionary scenarios using a likelihood ratio test (LRT). In parallel, we performed Tajima’s relative rate test (Tajima 1993) on each duplicate pair and NHP orthologous outgroup sequence. **Table 1** summarizes the total number of duplicate enhancer pairs in which one of the enhancers exhibits accelerated sequence evolution based on statistical significance in both tests (**Figure 3d**). In total, approximately 30% of all duplicate enhancer pairs using the rhesus macaque as an outgroup and 40% of all duplicate enhancer pairs with chimpanzee as an outgroup display signatures of sequence acceleration (**Table 1**). Hereafter, we will refer to the enhancer in a duplicate pair exhibiting significant acceleration as the “accelerating enhancer,” while the other enhancer will be called the “non-accelerating enhancer.”

We find that accelerating enhancers identified using the rhesus macaque as an outgroup are significantly less pleiotropic than their complementary non-accelerating enhancer (p < 0.002, paired sign test, **Figure 4a**, and **Supplementary Table 5**). Indeed, these enhancers were significantly more likely to be *entirely* tissue-specific (i.e., functioning as an enhancer in only one tissue with degree of pleiotropy = 1) than non-accelerating enhancers (odds ratio = 2.34, p < 0.003 Fisher’s Exact Test, **Supplementary Table 6**). These enhancers were also shorter and harbored fewer and less diverse transcription factor binding motifs than the non-accelerating enhancer in the pair (**Figure 4a**, and **Supplementary Table 5**). We found no significant differences in the genomic features of accelerating enhancers compared to non-accelerating enhancers when considering the most recent duplicates following the human-chimpanzee divergence (**Supplementary Table 7**), which may be due to the insufficient resolution of data at this time. Alternatively, they may reflect the instances of duplication events rather than the results of their evolutionary divergence. As such, for the following function annotation analyses, we will focus on the subset of duplicate enhancers exhibiting significant asymmetric evolution in the human-rhesus macaque comparison.

**Figure 4.**
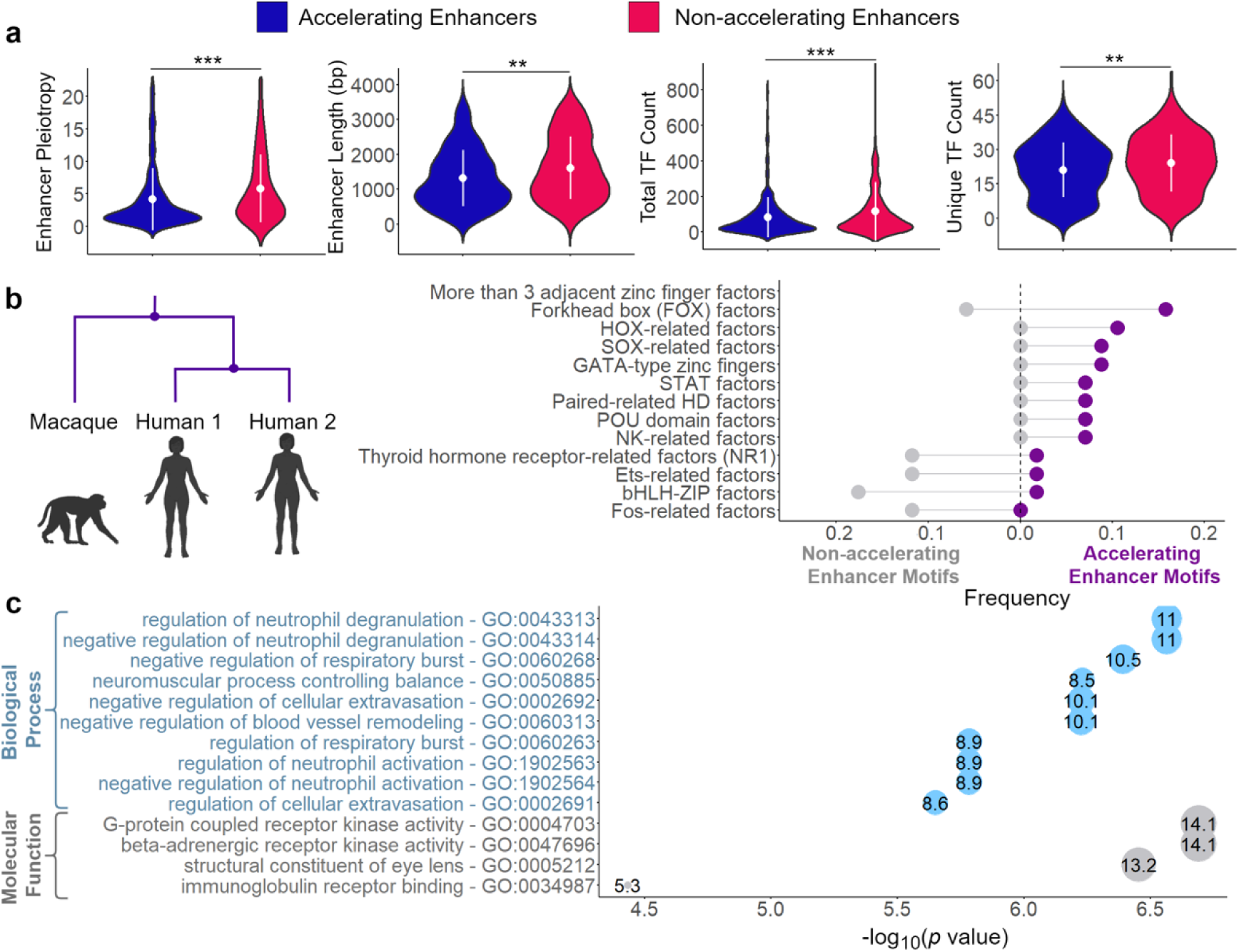
Features of duplicate enhancers exhibiting accelerated evolutions. **(a)** Violin plots comparing genomic attributes of duplicate enhancers experiencing accelerated sequence evolution compared to the associated non-accelerating enhancer. Reported p- p-values were calculated from paired two-sample sign tests (*** indicates p < 5.5 × 10^-5^ and ** indicates p < 9 × 10^-3^). (**b**) Frequency of transcription factor families from significantly and uniquely enriched TF motifs in either accelerating or non-accelerating duplicate enhancers. (**c**) Functional annotation of genes significantly associated with accelerating enhancers. In (**a-c**), accelerating enhancers were identified using the orthologous rhesus macaque region as the outgroup sequence.

We identified significantly enriched transcription factor (TF) binding motifs and associated transcription factor families within accelerating and non-accelerating enhancers using MEME (Bailey, et al. 2009) suite’s SEA software and the HOCOMOCO v11 core database of human TF binding motifs. Intriguingly, we show that enriched motifs found almost exclusively in accelerating enhancers compared to non-accelerating enhancers belong to FOX factors, HOX-related factors, and SOX-related factors, among others (**Figure 4b**).

Similarly, overrepresented motifs in non-accelerating enhancers are mostly associated with families relatively depleted in accelerating enhancers, including Fos-related factors, bHLH-ZIP factors, Ets-related factors, and thyroid hormone receptor-related factors (**Figure 4b**). Through Gene Ontology analysis using the Genomic Regions Enrichment of Annotations Tool (GREAT) v.3.0.0 (McLean, et al. 2010) and ShinyGO v0.741 (Ge, et al. 2020), we demonstrate that accelerating enhancers are significantly associated with genes enriched in immune functions and stress responses (FDR < 0.05 hypergeometric test, **Figure 4c** and **Supplementary Figure 2**). These analyses demonstrate that within duplicate pairs that emerged since human-rhesus macaque divergence, those exhibiting accelerated sequence evolution also exhibit functional attributes associated with tissue-specific evolution.

### The majority of accelerating duplicate enhancers gain novel tissue activity

Given the signature of asymmetrical sequence evolution within duplicate enhancer pairs, we endeavored to evaluate the corresponding effect on the collective breath of tissue activity of these enhancers. Specifically, we identified instances where accelerating enhancers in recently duplicated enhancer pairs gained activity in at least one novel tissue compared to the non-accelerating enhancer. These events may be indicative of regulatory neofunctionalization driven by a duplication event. We report that ∼75% of all accelerating enhancers consistently gained activity in novel tissues compared to the non-accelerating enhancer when considering asymmetrically evolving enhancers identified by either chimpanzee or rhesus macaque outgroup analysis (**Table 2**). This gain of enhancer regulatory function is primarily restricted to the addition of 1-2 novel tissues (**Figure 5a, c**).

**Figure 5.**
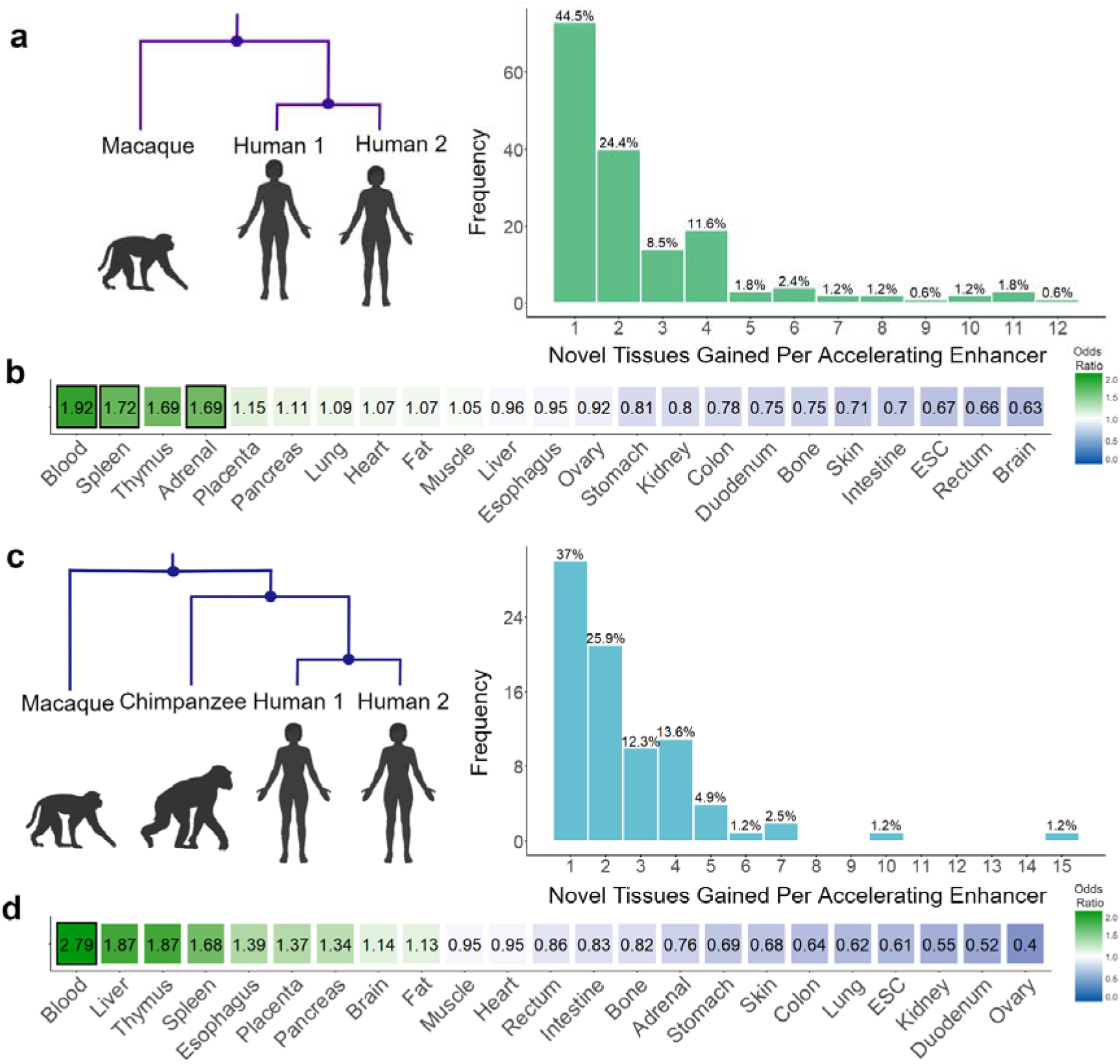
Gain of tissue activity and tissue enrichment of accelerating enhancers. In (**a** and **c**), the distribution of the number of novel tissues in which accelerating enhancers gain function compared to their non-accelerating mate for those identified using (**a**) rhesus macaque or (**c**) chimpanzee orthologous regions as outgroups. (**b** and **d**) The enrichment and significance of duplicate enhancers exhibiting accelerated evolution in each of the survey tissues compared to pleiotropy-matched enhancers in the corresponding tissue as the control background. Odds ratio and p-value are reported from Fisher’s Exact Test.

**Table 2.**
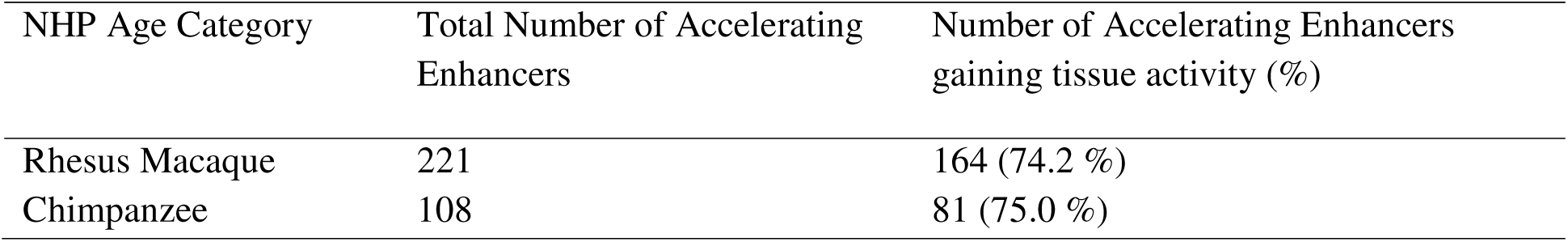
Gain of regulatory function in accelerating enhancers. Total number of accelerating enhancers that gain novel tissue activity compared to the tissue activities of their non-accelerating mate. Values are reported for accelerating enhancers identified using both non-human primate orthologous regions as outgroups.

We performed a permutation-based enrichment analysis of the occurrence of the accelerating enhancers in each tissue compared to the occurrence of pleiotropy-matched enhancers in the same tissue as the control background. In the subset of accelerating enhancers identified from the human-rhesus macaque comparison, there is a significant overrepresentation of blood, spleen, and adrenal enhancers (p < 0.05, Fisher’s Exact Test, **Figure 5b**). Although not significant, we note that brain enhancers showed the greatest depletion of these accelerating enhancers (Odds ratio = 0.63, p = 0.055, Fisher’s Exact Test). With respect to accelerating enhancers originating from duplication events following the more recent human-chimpanzee divergence, blood enhancers also show a significant enrichment compared to the control (OR = 2.79, p = 0.001, **Figure 4d**). In contrast, the brain enhancers show an enrichment for these more recently accelerating enhancers, although the enrichment is not significant (Odds ratio = 1.14, p = 0.77, Fisher’s Exact Test). Interestingly, when considering duplicates in the human-rhesus macaque comparison, we observe that non-accelerating enhancers (N=221 enhancers with mean pleiotropy = 5.84) in accelerating pairs are significantly more pleiotropic than those whose pairs are evolving symmetrically (N = 488 enhancers with mean pleiotropy = 3.94; p = 2.39 × 10^-5^ Mann–Whitney *U* test).

## Discussion

In this study, we explore the evolutionary origins of human enhancers through genomic duplication events. Many features of enhancer composition and organization may factor into the functional retention of duplicate enhancers. Enhancers have a reduced barrier of evolution due to their smaller size and ability to repurpose ancestral DNA and transposable elements lending themselves to greater instances of genesis through exaptation or spontaneous emergence (Rebeiz, et al. 2011; Villar, et al. 2015). In support of this evolutionary model, Fong and Capra (Fong and Capra 2021) reported that enhancers show enrichment for a “simple evolutionary architecture” where the underlying sequences came from a single rather than multiple evolutionary ages. Enhancers are also highly redundant in their interaction with genes, a feature that stabilizes the expression of target genes but may dilute the selective pressure on any single enhancer (Singh and Yi 2021). Importantly, in an analysis of 20 mammalian liver enhancers, Villar et al. (Villar, et al. 2015) reported that enhancer function experienced rapid evolutionary turnover.

Interestingly, some characteristics of duplicate enhancers appear to contribute to their evolutionary maintenance. We show that increased pleiotropy is a hallmark of duplicate enhancers. Additionally, these enhancers are longer, linked to a greater number of target genes to regulate, located closer to genic regions, and enriched for diverse transcription factor binding sites. Collectively, these features suggest that an increase in regulatory potential may play a factor in the retention of these enhancers in the genome. These trends are the most exaggerated when considering the “oldest” subset of duplicated enhancers (**Figure 2**).

Next, we sought to explore questions regarding the evolutionary trajectories and functionality of these duplicated regulatory regions. Specifically, how often does asymmetric evolution occur between recently duplicated pairs? In cases where one of the two duplicates exhibits accelerated evolution, how often do enhancers gain activity in novel tissues, indicating potential instances of duplication-driven regulatory neofunctionalization? Are there particular tissues that are overrepresented in accelerating enhancers? Using orthologous regions in rhesus macaque and chimpanzee genomes, we found that 30-40% of recently duplicated enhancer pairs experienced accelerated evolution in one enhancer copy. Notably, most of these accelerating enhancers (∼75%) gained novel tissue activity beyond the activity of their non-accelerating mate, suggestive of regulatory neofunctionalization. Consequently, the significant acceleration of these enhancers at the sequence level could be explained by positive selection associated with the gain of a novel tissue activity. Interestingly, the non-accelerating enhancers were among the most pleiotropic of all enhancers (mean pleiotropy = 5.84, **Supplementary Table 5**). Therefore, our observation is consistent with the idea that duplication of highly pleiotropic enhancers, which harbors a high degree of regulatory potential, contributes to the successful repurposing and subsequent selection to maintain novel function in the duplicated enhancer.

In evaluating the functional consequences of duplicate enhancers, we observe that all maintained duplicate enhancers, particularly accelerating enhancers, were significantly enriched in immune-related tissues including blood, spleen, and thymus samples. Moreover, accelerating enhancers were enriched in several transcription factor binding motifs that were previously associated with human-specific evolution of DNA methylation and chromatin accessibility (Caglayan, et al. 2023; Jeong, et al. 2021). In addition, the enriched target genes linked to accelerating enhancers correlated with stress and immune responses. These findings indicate that duplication of enhancers may have contributed to the evolution of novel immune-related functions in humans. We note that these observations are also concordant with previous results that the most recent enhancers experiencing positive selection were enriched for immune function (Moon, et al. 2019). The selection and retention of these enhancers may be partially driven by their function as many studies in diverse organisms have shown parallel and consistent signatures of positive selection in immune pathways (Barreiro and Quintana-Murci 2010; Kosiol, et al. 2008; Sackton, et al. 2007; Schlenke and Begun 2003).

Our study has several caveats. Systematically identifying enhancers from functional genomic data is still challenging, and we utilized mapping of specific histone marks to infer enhancer functionality across the genome. Even though it is widely used in recent studies (Roadmap Epigenomics, et al. 2015; Villar, et al. 2015), future advances in functional mapping studies may provide a more comprehensive list of enhancers. A substantial portion of the human genome consists of repetitive sequences, and many enhancers encode repetitive sequences (e.g., (Barth, et al. 2020; Rayan, et al. 2016)). Consequently, it is challenging to accurately map duplicate regions with high repeat contents (e.g., (Rhie, et al. 2021)). To enable sequence comparisons and confident identification of duplicate enhancers, here we used a cutoff of 50% repetitive sequences. Using a more stringent 70% cutoff yielded fewer duplicate enhancers, as expected. Nevertheless, the attribute and tissue differences between the duplicate enhancers and controls remained the same as those identified using a 50% cutoff (**Supplementary Figure 3**). As our understanding of the dynamics of repetitive sequences increases, future studies employing more sophisticated treatments of repetitive sequences could expand the scope of duplicate enhancers. Our results should be taken as a first look at conservatively identified duplicate enhancers.

## Methods

### Enhancer Dataset

The putative enhancer dataset and associated enhancer attributes were downloaded from https://github.com/soojinyilab/Enhancer_Dataset_2020. These attributes include number of tissues in which the enhancer was “present” based on enhancer chromatin state, the number of gene links per enhancer, the distances to the nearest gene measured in base pairs, and the number and diversity of TF motifs. The detailed methods of how these data were curated can be found in Singh and Yi (Singh and Yi 2021). Briefly, two representative samples were selected from the 127 epigenomes available from the NIH Roadmap Epigenomics Mapping Consortium (Roadmap Epigenomics Consortium, et al. 2015). These included 43 samples across 23 human tissues. Enhancer coordinates (state 6 and state 7) were obtained from the 15-state ChromHMM model and merged based on an overlap of 50% after outlier filtration. Corresponding TF motifs for each enhancer were identified using MEME suite’s FIMO package (Bailey, et al. 2009) and the HOCOMOCO v11 core database (Kulakovskiy, et al. 2016). Gene links were downloaded for the ENCODE+Roadmap dataset from the JEME repository (Cao, et al. 2017).

### Identification and Enrichment of Duplicate Enhancers

To identify duplicate enhancers from all putative enhancers, nucleotide sequences corresponding to the genomic coordinates of the original dataset of 646,419 enhancer regions were extracted from the human hg19 reference genome (GRCh37.p13). Following this extraction, RepeatMasker (Smit, et al. 2019) was run on the sequences to identify enhancers composed of highly repetitive elements (>50% repetitive content). To avoid spurious matches between highly repetitive regions in non-duplicate enhancers, these enhancers were excluded from the downstream analyses. Next, an all-by-all BLAST (*blastn* (Altschul, et al. 1990)) was performed on the remaining enhancer sequences with an e-value threshold of 1×10^-10^. Enhancer families were generated from the reciprocal BLAST hits where the two sequences overlapped by 50% the length of the shortest sequence. In total we report 2,362 enhancer families encompassing 7,118 enhancers.

For downstream sequence evolution analyses, it was critical to identify candidate pairs within duplicate enhancer families. To this end, multiple sequence alignments (MSAs) were generated for all enhancers within duplicate enhancer families using MAFFT (v7.310 (Katoh and Standley 2013)) the converted into evolutionary distance matrices for each family in MEGA (mega-cc v10 (Kumar, et al. 2018)). Evolutionary distances were measured by Kimura’s two-parameter (K2P) model (Kimura 1980). Following methodology similar to Park and Makova (Park and Makova 2009), single-linkage clustering of K2P distances was performed to merge enhancers within families into candidate duplicate pairs based on closet evolutionary distance.

For all duplicate enhancer enrichment analyses, 1,000 control datasets of length-matched non-duplicate enhancers were generated from the putative enhancer dataset. Enrichment p-values were reported as the ratio of the number of control values at least as extreme as the duplicate enhancer value over the total number of control datasets for each examined attribute. Fisher’s exact test and associate odds ratios were calculated to measure the enrichment or depletion of duplicate enhancers by examining the occurrence of duplicate enhancers within and without each tissue (observed value) compared to that of the control non-duplicate enhancer background (expected values).

### Identifying Asymmetric Evolution

To identify duplicate enhancers exhibiting asymmetric evolution, it was necessary to identify outgroups to measure sequence divergence. To that end, orthologous regions in two NHP genomes, rhesus macaque (rheMac10) and chimpanzee (panTro5), were identified using a reciprocal best BLAST hit (RBBH) approach (summarized in **Figure 3a**). For each enhancer in the duplicate enhancer dataset, a BLAST search was performed to find the “best hit” in each NPH genome. We then performed a reciprocal BLAST search of those “best hits” from the NHP genome against the human genome (GRCh37.p13). If the “best hit” for the reciprocal BLAST search returned the original enhancer region, the corresponding NHP was considered in the orthologous region. Notable, for recent duplicates resulting from events following the divergence of the human-NPH lineage, it is expected that there would be a shared “best hit” in the NPH genome as there would be a single-copy orthologous region. Consequently, the reciprocal “best hit” from the NPH region to the human genome would match the one of the two recent duplicates (presumably the duplicate exhibiting less sequence divergence from the NPH genome). To identify this subset, duplicate enhancer pairs that shared “best hits” in the NPH genome and did not map to multiple NPH regions were retained as candidate recent duplicates. These regions were classified to have single-copy orthologous regions in one or both NPH genomes (**Figure 3b**).

Utilizing these single-copy orthologous sequences, baseml from PAML (Yang 2007) was performed to identify instances of asymmetric evolution. Specifically, for each recently duplicated enhancer pair, a phylogenetic tree was generated from a multiple sequence alignment (MSA) of the two enhancers and the outgroup primate orthologous region and then tested to better fit a molecular clock model in which branches have the same rate or a free-rates clock model in which the rates of branches can vary. The free-rate model accounts for cases of asymmetric evolution of duplicated enhancer pairs. Baseml reports the log likelihoods of each model which were then compared by a likelihood ratio test to determine duplicate pairs whose divergence was significantly better fit by the free-rate model. In tandem, the MSAs were analyzed by Tajima’s relative rate test (Tajima 1993) implemented in MEGA (mega-cc v10 (Kumar, et al. 2018)) to confirm or reject the molecular clock hypothesis indicative of symmetric sequence evolution based on the sequence divergence of the duplicate pair. Duplicate pairs with significant signatures of asymmetric evolution in both analyses were included in subsequent analyses.

### Functional Annotation of Accelerating Enhancers

Functional annotation of TF motifs and associated TF families for duplicate enhancer exhibiting asymmetric evolution was performed by MEME suite’s (Bailey, et al. 2009) SEA package and the HOCOMOCO v11 core database of human TF binding motifs. First, the enrichment of TF motifs was determined for both accelerating and non-accelerating enhancers independently using length-matched control regions as the genomic background. The unique TF motifs in each enrichment set were then identified as the motifs unique enrichment in accelerating enhancers compared to their non-accelerating enhancer mate. From these enrichments, the frequency of associated TF families was reported. Gene ontology of the target genes of accelerating enhancers was performed independently using two tools. The Genomic Regions Enrichment of Annotation Tool (GREAT v3.0.0. (McLean, et al. 2010)), is specifically designed to annotate biological meaning to genes linked to non-coding *cis*-regulatory elements based on proximity. The associated significant GO terms related to Biological Process and Molecular Function were identified for accelerating enhancers using all duplicate enhancers as the background set and a FDR threshold > 0.05. Independently, the JEME gene-links annotated to enhancers in Singh and Yi (Singh and Yi 2021), were extracted for the subset of enhancers exhibiting accelerated sequence evolution. These genes were tested for significant enrichment in ShinyGO v0.741 (Ge, et al. 2019) against a background set of all duplicate enhancer linked genes in the total duplicate enhancer dataset. The higher GO terms are reported for the enriched genes.

## Notes

### Competing Interest Statement

The authors have declared no competing interest.

## References

Abascal F, et al. 2020. Expanded encyclopaedias of DNA elements in the human and mouse genomes. Nature 583: 699–710. doi: 10.1038/s41586-020-2493-4

Acharya D, Ghosh TC 2016. Global analysis of human duplicated genes reveals the relative importance of whole-genome duplicates originated in the early vertebrate evolution. BMC Genomics 17: 71. doi: 10.1186/s12864-016-2392-0

Allan CM, Walker D, Taylor JM 1995. Evolutionary Duplication of a Hepatic Control Region in the Human Apolipoprotein E Gene Locus: IDENTIFICATION OF A SECOND REGION THAT CONFERS HIGH LEVEL AND LIVER-SPECIFIC EXPRESSION OF THE HUMAN APOLIPOPROTEIN E GENE IN TRANSGENIC MICE (&#x2217;). Journal of Biological Chemistry 270: 26278–26281. doi: 10.1074/jbc.270.44.26278

Altschul DA, Gish W, Miller W, Myers EW, Lipman DJ 1990. Basic local alignment search tool. J. Mol. Biol. 215: 403–410.

Bailey TL, et al. 2009. MEME Suite: tools for motif discovery and searching. Nucleic Acids Research 37: W202–W208.

Banerji J, Rusconi S, Schaffner W 1981. Expression of a β-globin gene is enhanced by remote SV40 DNA sequences. Cell 27: 299–308.

Barreiro LB, Quintana-Murci L 2010. From evolutionary genetics to human immunology: how selection shapes host defence genes. Nature Reviews Genetics 11: 17–30. doi: 10.1038/nrg2698

Barth NKH, Li L, Taher L 2020. Independent Transposon Exaptation Is a Widespread Mechanism of Redundant Enhancer Evolution in the Mammalian Genome. Genome Biology and Evolution 12: 1–17. doi: 10.1093/gbe/evaa004

Caglayan E, et al. 2023. Molecular features driving cellular complexity of human brain evolution. Nature 620: 145–153. doi: 10.1038/s41586-023-06338-4

Cao Q, et al. 2017. Reconstruction of enhancer–target networks in 935 samples of human primary cells, tissues and cell lines. Nature Genetics 49: 1428–1436. doi: 10.1038/ng.3950

Croft B, et al. 2018. Human sex reversal is caused by duplication or deletion of core enhancers upstream of SOX9. Nature Communications 9: 5319. doi: 10.1038/s41467-018-07784-9

ENCODE_consortium 2012. An integrated encyclopedia of DNA elements in the human genome. Nature 489: 57–74.

Fong SL, Capra JA 2021. Modeling the Evolutionary Architectures of Transcribed Human Enhancer Sequences Reveals Distinct Origins, Functions, and Associations with Human Trait Variation. Molecular Biology and Evolution 38: 3681–3696. doi: 10.1093/molbev/msab138

Force A, et al. 1999. Preservation of duplicate genes by complementary, degenerative mutations. Genetics 151: 1531–1545.

Ge SX, Jung D, Yao R 2020. ShinyGO: a graphical gene-set enrichment tool for animals and plants. Bioinformatics 36: 2628–2629. doi: 10.1093/bioinformatics/btz931

Goode DK, Callaway HA, Cerda GA, Lewis KE, Elgar G 2011. Minor change, major difference: divergent functions of highly conserved cis-regulatory elements subsequent to whole genome duplication events. Development 138: 879–884. doi: 10.1242/dev.055996

Huh I, Mendizabal I, Park T, Yi SV 2018. Functional conservation of sequence determinants at rapidly evolving regulatory regions across mammals. PLoS Computational Biology 14: e1006451. doi: 10.1371/journal.pcbi.1006451

Innan H, Kondrashov FA 2010. The evolution of gene duplications: classifying and distinguishing between models. Nature Reviews Genetics 11: 97–108.

Jeong H, et al. 2021. Evolution of DNA methylation in the human brain. Nature Communications 12: 2021. doi: 10.1038/s41467-021-21917-7

Katoh K, Standley DM 2013. MAFFT Multiple Sequence Alignment Software Version 7: Improvements in Performance and Usability. Molecular Biology and Evolution 30: 772–780. doi: 10.1093/molbev/mst010

Keller TE, Yi SV 2014. DNA methylation and evolution of duplicate genes. Proceedings of the National Academy of Sciences in press.

Kimura M 1980. A simple method for estimating evolutionary rate of base substitution through comparative studies of nucleotide sequences. J. Mol. Evol. 16: 111–120.

Kosiol C, et al. 2008. Patterns of Positive Selection in Six Mammalian Genomes. PLoS Genet 4: e1000144. doi: 10.1371/journal.pgen.1000144

Kulakovskiy IV, et al. 2016. HOCOMOCO: expansion and enhancement of the collection of transcription factor binding sites models. Nucleic Acids Research 44: D116–D125. doi: 10.1093/nar/gkv1249

Kumar S, Stecher G, Li M, Knyaz C, Tamura K 2018. MEGA X: Molecular Evolutionary Genetics Analysis across Computing Platforms. Molecular Biology and Evolution 35: 1547–1549. doi: 10.1093/molbev/msy096

Kvon EZ, Waymack R, Gad M, Wunderlich Z 2021. Enhancer redundancy in development and disease. Nature Reviews Genetics 22: 324–336. doi: 10.1038/s41576-020-00311-x

Lettice LA, et al. 2014. Development of five digits is controlled by a bipartite long-range cis-regulator. Development 141: 1715–1725. doi: 10.1242/dev.095430

Li W-H, Gu Z, Wang H, Nekrutenko A 2001. Evolutionary analyses of the human genome. Nature 409: 847–849. doi: 10.1038/35057039

Long HK, Prescott SL, Wysocka J 2016. Ever-Changing Landscapes: Transcriptional Enhancers in Development and Evolution. Cell 167: 1170–1187. doi: 10.1016/j.cell.2016.09.018

Lynch M, Conery JS 2000. The evolutionary fate and consequences of duplicate genes. Science 290: 1151–1155.

Lynch M, Force A 2000. The probability of duplicate gene preservation by subfunctionalization. Genetics 154: 459–473.

McLean CY, et al. 2010. GREAT improves functional interpretation of cis-regulatory regions. Nature biotechnology 28: 495–501. doi: 10.1038/nbt.1630

Moon JM, Capra JA, Abbot P, Rokas A 2019. Signatures of Recent Positive Selection in Enhancers Across 41 Human Tissues. G3 Genes|Genomes|Genetics 9: 2761-2774. doi: 10.1534/g3.119.400186

Ngcungcu T, et al. 2017. Duplicated Enhancer Region Increases Expression of CTSB and Segregates with Keratolytic Winter Erythema in South African and Norwegian Families. The American Journal of Human Genetics 100: 737–750. doi: 10.1016/j.ajhg.2017.03.012

Ohno S. 1970. Evolution by Gene Duplication. Berlin: Springer-Verlag.

Osterwalder M, et al. 2018. Enhancer redundancy provides phenotypic robustness in mammalian development. Nature 554: 239–243. doi: 10.1038/nature25461

Park C, Makova KD 2009. Coding region structural heterogeneity and turnover of transcription start sites contribute to divergence in expression between duplicate genes. Genome Biology 10: R10. doi: 10.1186/gb-2009-10-1-r10

Rayan NA, del Rosario RCH, Prabhakar S 2016. Massive contribution of transposable elements to mammalian regulatory sequences. Seminars in Cell & Developmental Biology 57: 51-56. doi: 10.1016/j.semcdb.2016.05.004

Rebeiz M, Jikomes N, Kassner VA, Carroll SB 2011. Evolutionary origin of a novel gene expression pattern through co-option of the latent activities of existing regulatory sequences. Proceedings of the National Academy of Sciences 108: 10036–10043. doi: 10.1073/pnas.1105937108

Rhie A, et al. 2021. Towards complete and error-free genome assemblies of all vertebrate species. Nature 592: 737–746. doi: 10.1038/s41586-021-03451-0

Roadmap Epigenomics C, et al. 2015. Integrative analysis of 111 reference human epigenomes. Nature 518: 317–330. doi: 10.1038/nature14248

Sabarís G, Laiker I, Preger-Ben Noon E, Frankel N 2019. Actors with Multiple Roles: Pleiotropic Enhancers and the Paradigm of Enhancer Modularity. Trends In Genetics 35: 423–433.

Sackton TB, et al. 2007. Dynamic evolution of the innate immune system in Drosophila. Nature Genetics 39: 1461–1468. doi: 10.1038/ng.2007.60

Schlenke TA, Begun DJ 2003. Natural Selection Drives Drosophila Immune System Evolution. Genetics 164: 1471–1480. doi: 10.1093/genetics/164.4.1471

Si N, et al. 2020. Duplications involving the long range HMX1 enhancer are associated with human isolated bilateral concha-type microtia. Journal of Translational Medicine 18: 244. doi: 10.1186/s12967-020-02409-6

Singh D, Yi SV 2021. Enhancer Pleiotropy, Gene Expression, and the Architecture of Human Enhancer–Gene Interactions. Molecular Biology and Evolution 38: 3898–3909. doi: 10.1093/molbev/msab085

Smit AFA, Hubely R, Green P 2019. RepeatMasker Open-4.0.

Tajima F 1993. Simple methods for testing the molecular evolutionary clock hypothesis. Genetics 135: 599–1465.

Villar D, et al. 2015. Enhancer Evolution across 20 Mammalian Species. Cell 160: 554–566.

Waymack R, Fletcher A, Enciso G, Wunderlich Z 2020. Shadow enhancers can suppress input transcription factor noise through distinct regulatory logic. eLife 9: e59351. doi: 10.7554/eLife.59351

Yang Z 2007. PAML 4: Phylogenetic Analysis by Maximum Likelihood. Molecular Biology and Evolution 24: 1586–1591. doi: 10.1093/molbev/msm088

