## Supplementary Files for "Evolution of Enhancers through Duplication"

### Supplementary Material

**Supplementary Figure 1. Distribution of K2P distances between duplicate enhancer pairs. The d**ashed line denotes the mean K2P value of the dataset.

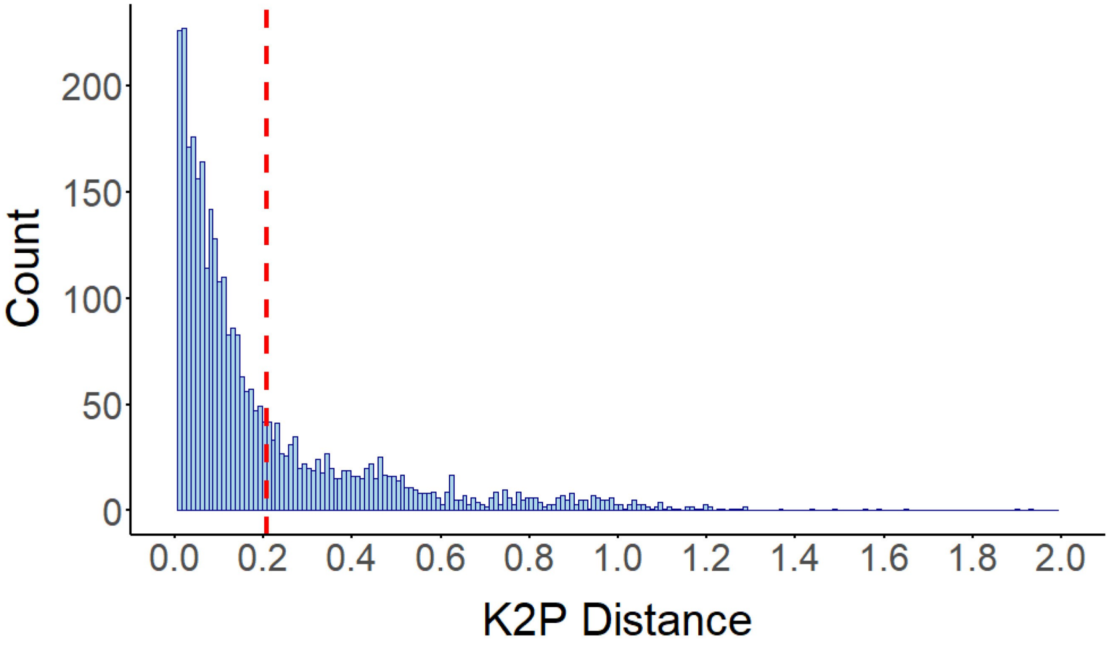

**Supplementary Figure 2. Gene Ontology of genes associated with accelerating duplicate enhancers.** Higher GO terms for Biological Process and Molecular Function annotation for genes enriched in the accelerating duplicate enhancer dataset identified using either the (a) rhesus macaque or (b) chimpanzee orthologous regions as outgroups.

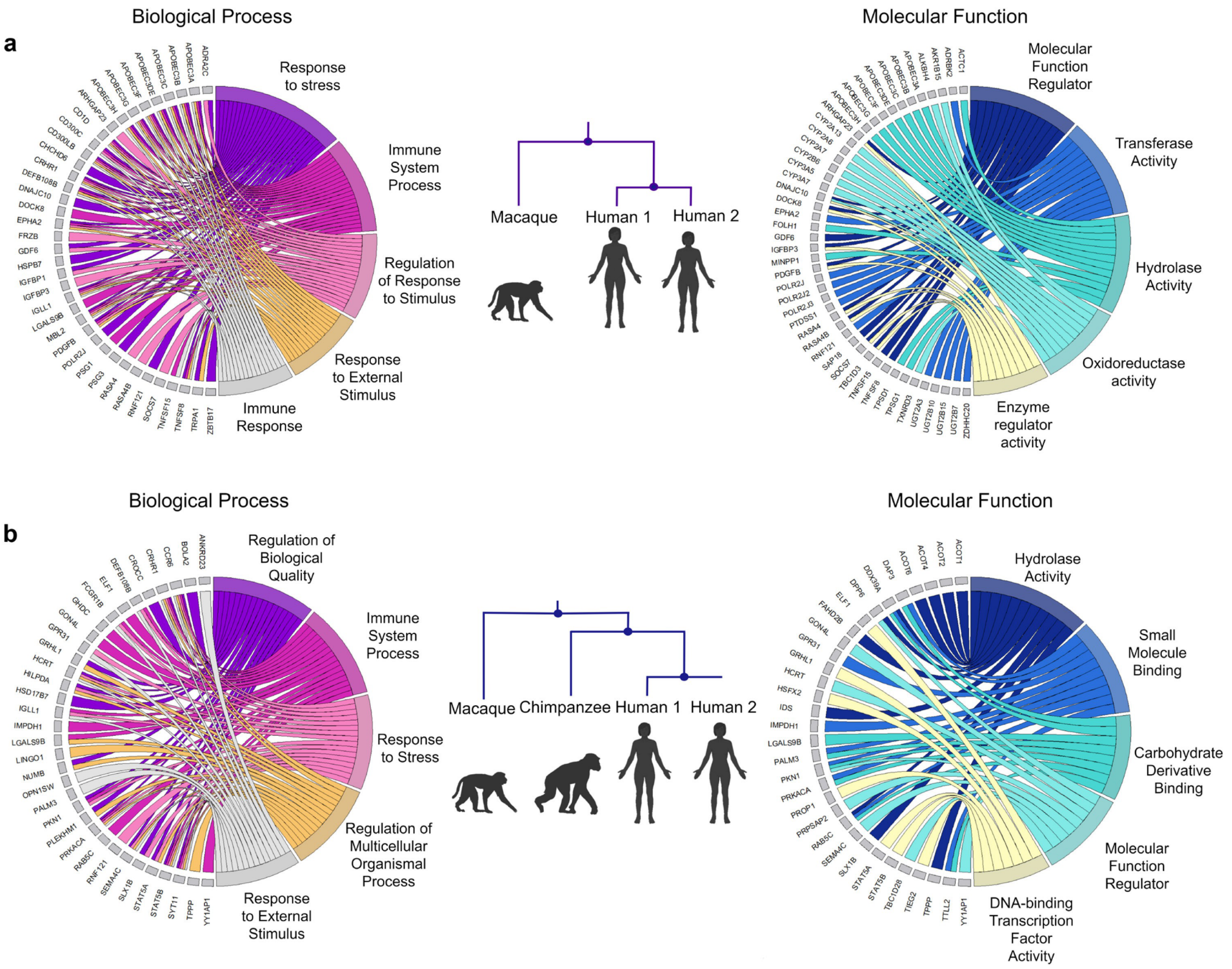

**Supplementary Figure 3. Genomic characteristics of duplicate enhancers. (a)** Enrichment of six genomic attributes of duplicate enhancers are shown for duplicate enhancers defined under different criteria. For all attributes (a-f), p < 0.001 (illustrated as *) based on 1,000 bootstraps. Error bars indicate standard deviation. (**b**) Enrichment of duplicate enhancers in all surveyed tissues compared to length-matched non-duplicate control enhancers. Odds ratio and p-value are reported from Fisher's Exact Test considering the occurrence of duplicate enhancers active or not active in each tissue compared to the expected pattern from the control enhancers.

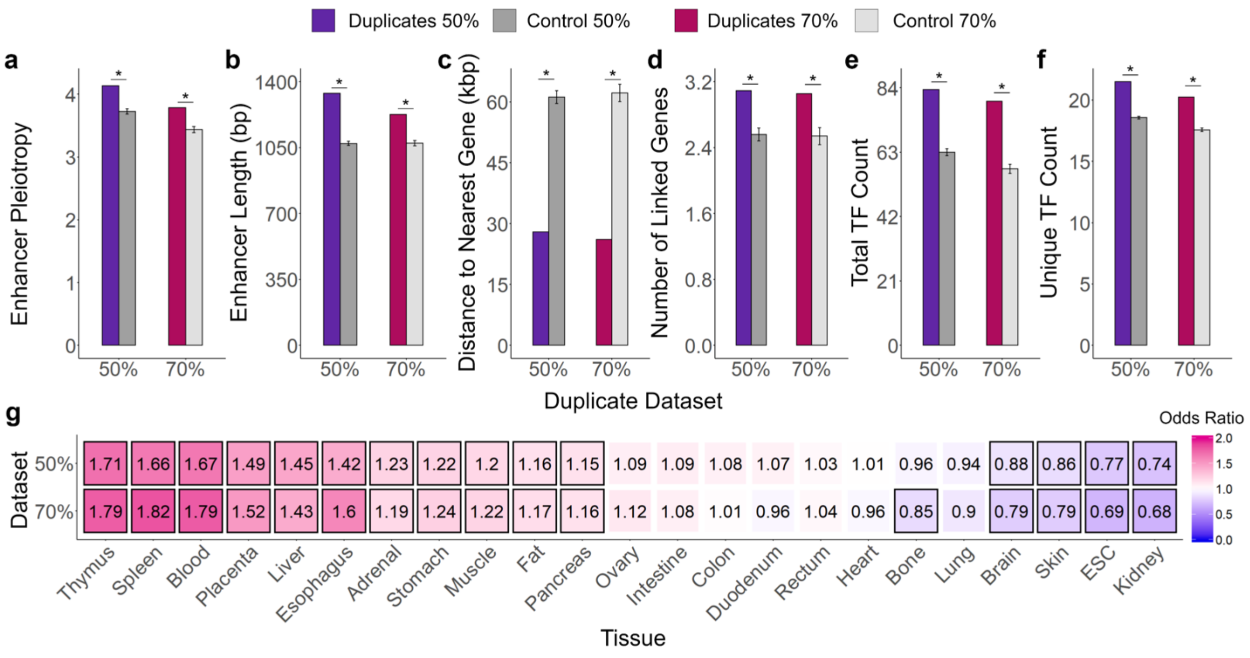

**Supplementary Table 1.** Mean attribute values for duplicate enhancers compared length-matched non-duplicate enhancer controls. The reported p-values as based on 1,000 bootstraps of the control regions.

| Enhancer Attribute | Duplicate Enhancers | Control Enhancers | P-value |
|---|---|---|---|
| Pleiotropy | 4.14 ± 4.4 | 3.72 ± 0.04 | < 0.001 |
| Length | 1338.0 ± 925.4 | 1072.3 ± 10.6 | < 0.001 |
| Distance to Nearest Gene (kbp) | 28.0 ± 72.3 | 61.2 ± 1.6 | < 0.001 |
| Number of Target Gene Links | 3.1 ± 2.8 | 2.6 ± 0.08 | < 0.001 |
| Total TF Binding Motifs | 83.5 ± 110.5 | 63.0 ± 1.1 | < 0.001 |
| Unique TF Binding Motifs | 21.5 ± 12.2 | 18.6 ± 0.11 | < 0.001 |

**Supplementary Table 2 Enrichment of duplicate enhancers across all surveyed tissues compared to length-matched non-duplicate control enhancers.** Odds ratio and p-value are reported from Fisher's Exact Test considering the occurrence of duplicate enhancers active or not active in each tissue compared to the expected pattern from the control enhancers.

| Tissue | Odds Ratio | P-value |
|---|---|---|
| ESC | 0.7717 | $7.06 \times 10^{-9}$ |
| Blood | 1.668 | $6.99 \times 10^{-34}$ |
| Skin | 0.8586 | $5.01 \times 10^{-4}$ |
| Fat | 1.157 | 0.003 |
| Liver | 1.4535 | $9.48 \times 10^{-15}$ |
| Brain | 0.8830 | 0.002 |
| Colon | 1.078 | 0.133 |
| Duodenum | 1.074 | 0.279 |
| Esophagus | 1.422 | $8.16 \times 10^{-10}$ |
| Adrenal | 1.226 | $1.65 \times 10^{-5}$ |
| Heart | 1.005 | 0.918 |
| Intestine | 1.091 | 0.058 |
| Kidney | 0.7446 | $8.39 \times 10^{-6}$ |
| Lung | 0.9430 | 0.146 |
| Muscle | 1.203 | $4.71 \times 10^{-6}$ |
| Placenta | 1.486 | $1.42 \times 10^{-19}$ |
| Stomach | 1.223 | $1.43 \times 10^{-6}$ |
| Thymus | 1.705 | $6.96 \times 10^{-27}$ |
| Ovary | 1.087 | 0.137 |
| Pancreas | 1.152 | 0.001 |
| Rectum | 1.034 | 0.533 |
| Spleen | 1.663 | $3.72 \times 10^{-24}$ |
| Bone | 0.9562 | 0.369 |

**Supplementary Table 3. Mean attribute values of duplicate enhancers binned evenly by K2P distance between duplicate pairs compared to the mean value of 1,000 bootstraps of the control non-duplicate enhancers.** The K2P ranges within each bin are as follows: Bin 1 K2P = 0-0.33, Bin 2 K2P = 0.33-0.67, Bin 3 K2P = 0.67-1.00, and Bin 4 K2p > 1.

| Enhancer Attribute | Control Mean | Bin 1 Mean | Bin 2 Mean | Bin 3 Mean | Bin 4 Mean |
|---|---|---|---|---|---|
| Pleiotropy | 3.72 ± 0.04 | 3.84 ± 4.2 | 4.5 ± 4.7 | 6.8 ± 5.1 | 7.7 ± 5.5 |
| Length (bp) | 1072.3 ± 10.6 | 1215.4 ± 865.3 | 1524.7 ± 964.2 | 2345.8 ± 739.5 | 2555.2 ± 855.9 |
| Total TF Count | 63.0 ± 1.1 | 77.6 ± 106.6 | 86.2 ± 98.5 | 140.1 ± 154.4 | 153.8 ± 140.7 |
| Unique TF Count | 18.6 ± 0.11 | 20.2 ± 12.0 | 23.7 ± 11.5 | 31.3 ± 9.8 | 32.3 ± 10.8 |

**Supplementary Table 4. Total counts (and percent) of duplicate enhancers with variations in copy number in the human, chimpanzee, and rhesus macaque genomes.** Each column is labeled by the copy number of the enhancer found in the human- chimpanzee-rhesus macaque genomes respectively. The (?) symbol represents instances where the human enhancer did not map an orthologous region of the corresponding non-human primate genome. Enhancers in the 2-1-2 column represent examples of duplication 'loss' in the chimpanzee genome.

| Human-Chimpanzee-Macaque (H-C-M) Duplicate Enhancer Copy Number (%) | | | | | |
|---|---|---|---|---|---|
| 2-1-1 | 2-2-1 | 2-1-2 | 2-?-2 | 2-2-2 | Total |
| 260 (10.6%) | 444 (18%) | 69 (2.8%) | 34 (1.4%) | 1654 (67.2%) | 2461 (100%) |

**Supplementary Table 5. Mean attribute values for duplicate enhancers exhibiting accelerated evolution compared to their non-accelerating mate.** Accelerating enhancers were identified using the orthologous region in the rhesus macaque genome as an outgroup. Reported p-values were calculated from paired two-sample sign tests.

| Enhancer Attribute | Mean Accelerating Enhancer | Mean Non-accelerating Enhancer | p-value |
|---|---|---|---|
| Pleiotropy | 4.18 | 5.84 | $5.90 \times 10^{-7}$ |
| Length (bp) | 1314.9 | 1613.6 | $8.72 \times 10^{-3}$ |
| Total TF Count | 58.6 | 71.5 | $5.42 \times 10^{-5}$ |
| Unique TF Count | 21.8 | 24.4 | $1.48 \times 10^{-3}$ |

**Supplementary Table 6. Values for a contingency table used to identify the enrichment of accelerating enhancers as *entirely* tissue-specific (pleiotropy = 1) compared to their corresponding non-accelerating enhancers.** Accelerating enhancers identified using both non-human primate orthologous regions as outgroups are reporting. Odds ratio and p-value are reported from Fisher's Exact Test.

| NHP Age Category | Accelerating Enhancer Pleiotropy = 1 | Accelerating Enhancer Pleiotropy > 1 | Non-accelerating Enhancer Pleiotropy = 1 | Non-accelerating Enhancer Pleiotropy > 1 | Odds Ratio (*p*-value) |
|---|---|---|---|---|---|
| Rhesus Macaque | 87 | 134 | 48 | 173 | 2.34 (**0.0001**) |
| Chimp | 65 | 135 | 57 | 143 | 1.21 (0.44) |

**Supplementary Table 7. Mean attribute values for duplicate enhancers exhibiting accelerated evolution compared to their non-accelerating mate.** Accelerating enhancers were identified using the orthologous region in the chimpanzee genome as an outgroup. Reported p-values were calculated from paired two-sample sign tests.

| Enhancer Attribute | n Accelerating Enhancer | Mean Non-accelerating Enhancer | Sign Test p-value |
|---|---|---|---|
| Pleiotropy | 4.50 | 4.69 | 0.51 |
| Length (bp) | 1472.2 | 1398.1 | 0.13 |
| Total TF Count | 104.8 | 90.4 | 0.84 |
| Unique TF Count | 24.2 | 23.6 | 0.25 |

**Supplementary Table 8.** Rate of enhancer duplications calculated using total number of duplicates identified with single-copy orthologous regions in both non- human primate genomes.

| NHP Age Category | Number of duplicate enhancers | Time since Human Divergence (my) | Enhancer Duplication Rate (duplications/enhancer/million years) |
|---|---|---|---|
| Rhesus Macaque | 738 | 25 | $4.57 \times 10^{-5}$ |
| Chimpanzee | 260 | 7 | $5.73 \times 10^{-5}$ |
